# Multiple rod layers increase the speed and sensitivity of vision in nocturnal reef fishes

**DOI:** 10.1101/2022.11.27.518067

**Authors:** Lily G. Fogg, Wen-Sung Chung, Fabio Cortesi, N. Justin Marshall, Fanny de Busserolles

## Abstract

Multibank retinas have rod photoreceptors stacked into multiple layers. They are found in many species of fish that inhabit dim environments and are one of the most common visual adaptations in the deep-sea. Despite its prevalence, the function of multibank retinas remained unknown. Two predominant theories, neither of which has been tested, have emerged: 1) they enhance sensitivity in dim light, and 2) they allow colour vision in dim light. To investigate the sensitivity hypothesis, we performed electrophysiological recordings and compared the rod pigments of three species of nocturnal reef fishes, two with a multibank retina (*Neoniphon sammara* and *Myripristis violacea*) and a control species with a single rod bank (*Ostorhinchus compressus*). Results indicated that nocturnal reef fishes with a multibank retina have higher temporal resolution of vision, as indicated by electrophysiology, and that their rhodopsin proteins likely also have faster retinal release kinetics, as suggested by amino acid substitutions. Electrophysiology also showed that the multibank retina conferred greater sensitivity to both dim and bright intensities than a single rod bank and this occurred at times when rod-derived signals usually dominate the visual response. This study provides the first functional evidence for enhanced dim-light sensitivity using a multibank retina while also suggesting novel roles for the adaptation in enhancing bright-light sensitivity and the speed of vision.

**Significance:** Most vertebrates have one layer of the dim-light active rod photoreceptors; however, some species have multiple layers, known as a multibank retina. We used electrophysiology on nocturnal reef fishes with and without multibank retinas to evaluate the sensory advantage of having multiple rod layers. We show that fish with multibank retinas have both faster vision and enhanced sensitivity to bright and dim light intensities. Thus, we resolve for the first time the function of multibank retinas – one of the most common visual adaptations in the deep sea. Our findings highlight an unconventional vertebrate visual system as well as the visual capabilities of fishes from the most vast (deep sea) and vibrant (reefs) ecosystems on the planet.

## Introduction

A great diversity of visual adaptations has evolved across the animal kingdom to permit vision in a myriad of ecological niches. For example, in invertebrates, these visual adaptations range from the 12 colour photoreceptors of the mantis shrimp (1) to the polarisation vision of locusts used for celestial navigation (2). While in terrestrial vertebrates, these adaptations include the hybrid cone-like rods of colubrid snakes (3, 4) to the highly sensitive eyes of some geckos that can discriminate colour in moonlight (5). Marine fishes are no exception to this diversity (6). To catch as many photons as possible, marine fishes living in dim-light environments such as in the deep-sea or at night show arguably the most extreme visual adaptations among vertebrates (7, 8). These scotopic adaptations include enlarged eyes or tubular eye structures (9, 10), high expression of the rod opsin gene, *rh1* (11, 12), high rod densities (13, 14), and thick photoreceptor layers, either through longer rods or multiple layers of rods, known as a multibank retina (12, 15, 16). Although many of these adaptations have been attributed to increasing sensitivity, the function of the multibank retina remained untested.

Multibank retinas consist of 2-28 layers of stacked rods (16, 17) and have been found in representatives from at least 38 teleost fish families (7, 18), the vast majority of which are deep-sea species (7). Two predominant theories have been suggested to explain their function. The first theory proposes that multibank retinas enhance luminous sensitivity by increasing the cumulative rod outer segment length available for photon capture (19). The second theory suggests that they allow colour vision in dim light through spectral filtering at each layer and an opponent comparison between the layers (20). Until now, few studies have examined the function of multibank retinas (21-23), due to the difficulty in accessing, handling, and maintaining deep-sea fishes (16, 24). However, the recent characterisation of multibank retinas in an easily accessible family of nocturnal coral reef fishes, Holocentridae (12), enabled us to test the sensitivity hypothesis.

Holocentridae is composed of two sub-families: squirrelfishes (Holocentrinae) and soldierfishes (Myripristinae). They mainly inhabit shallow depth ranges, however, a few species dwell as deep as 640 metres (25, 26). Holocentrids are nocturnal (27) and as such, they have a typical dim light-adapted visual system with large eyes (9), a rod-dominated retina (12, 28), a short focal length (15), a high summation of rods onto ganglion cells (GC) (29) and *rh1* genes with spectral sensitivities that are tuned to the dominant wavelengths at their prevalent depth (30). They also possess a highly developed multibank retina, with up to 7 and 17 banks in squirrelfishes and soldierfishes, respectively (12). However, holocentrids have also retained some photopic adaptations, including the potential for dichromatic colour vision (12).

In this study, the sensitivity theory was tested by assessing the visual systems of two holocentrid species (*Neoniphon sammara* and *Myripristis violacea*), and a non-multibank control species (*Ostorhinchus compressus*). Firstly, retinal structure was examined using histology. Then, the luminous sensitivity and temporal resolution of their vision was studied by recording the electrophysiological response of the whole eye to different light stimuli, a technique known as electroretinography (ERG) (31-34). Finally, we estimated the retinal release rate of the rhodopsin paralog expressed in the rods of each species. Overall, this study sheds light on the unresolved function of a prevalent but understudied visual adaptation in the deep sea as well as offering a broader insight into the vision of nocturnal reef fishes.

## Results

### Holocentrids have high rod densities and high scotopic summation

Retinal architecture and cell densities were assessed in *O. compressus, N. sammara* and *M. violacea* (*n*=1). All three species had duplex retinas composed of both rods and cones. However, while *O. compressus* only had a single layer of rods (Fig. 1Ai, Fig. S1), *N. sammara* and *M. violacea* had a maximum of 6 and 14 banks of rods, respectively (Fig. 1Aii-iii, Fig. S1). The densities of all cell types were heterogeneous across the retina in all species (Fig. 1B, Table S2). In every region, the highest rod densities and summation of rods onto GC occurred in *M. violacea* (peak rod densities, 21,296 cells/0.01mm^2^; peak rod:GC ratio, 1,651.5 rods/GC) followed by *N. sammara* (peak rod, 12,403 cells/0.01mm^2^; peak rod:GC, 332.6 rods/GC) and then *O. compressus* (peak rod, 3,545 cells/0.01mm^2^; peak rod:GC, 78.1 rods/GC). An inverse pattern was observed for cone and GC densities in all regions, with *O. compressus* having the highest densities and *M. violacea* the lowest (*O. compressus*: 72.8 cells/0.01mm^2^ and 97.5 cells/0.01mm^2^ for peak cone and GC, respectively; *N. sammara*: 49.4 cells/0.01mm^2^ and 71.0 cells/0.01mm^2^; *M. violacea*: 19.5 cells/0.01mm^2^ and 29.2 cells/0.01mm^2^). Finally, inner nuclear layer (INL) cell densities were also highest in *O. compressus* and lowest in *M. violacea* for most regions (*i*.*e*., dorsal, central and temporal) (peak INL, *O. compressus*: 1108 cells/0.01mm^2^; *N. sammara*: 789 cells/0.01mm^2^; *M. violacea*: 638 cells/0.01mm^2^).

**Fig. 1.**
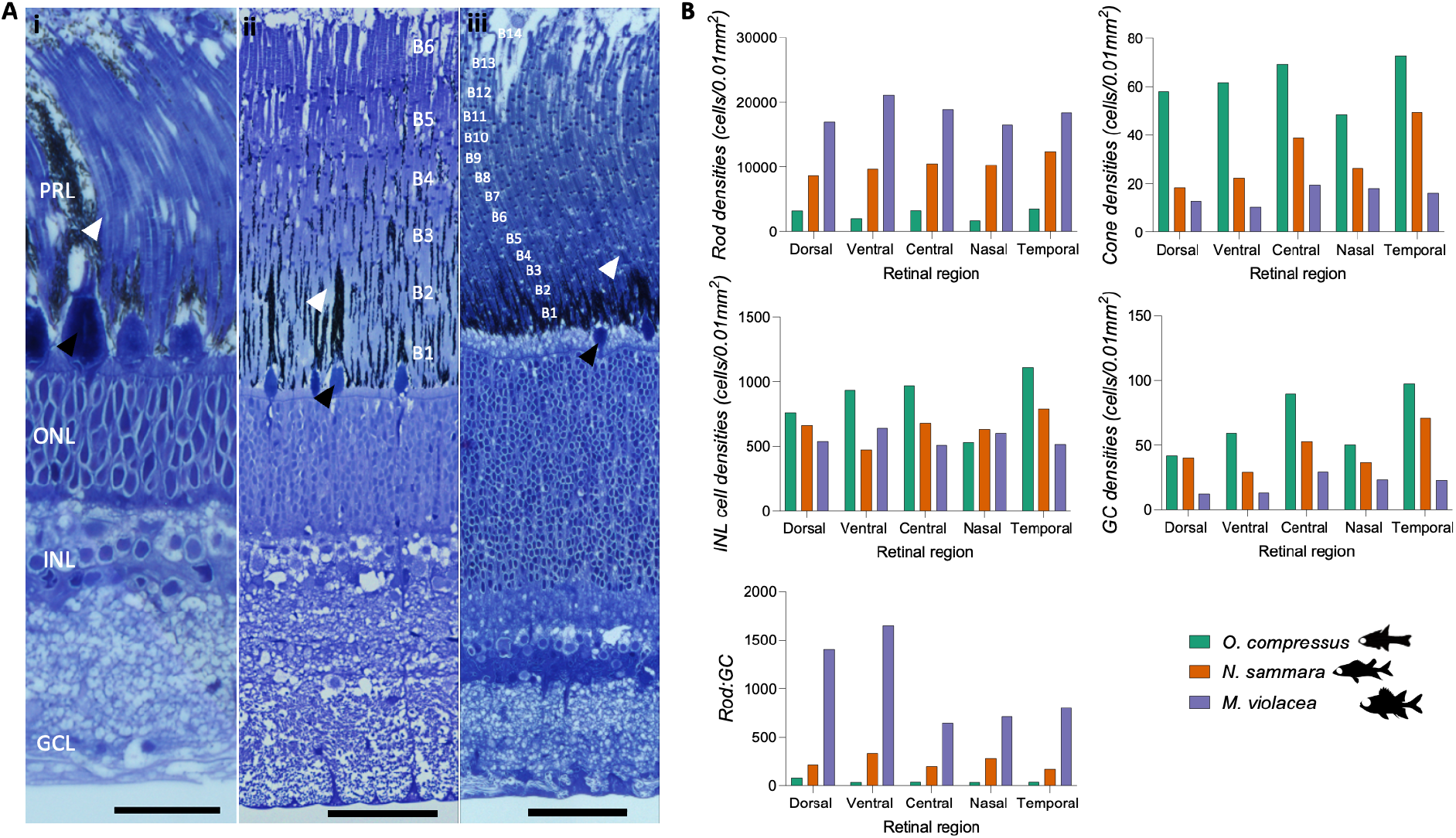
Retinal structure and cell densities. **A**. Representative radial sections from the retina of **i**) *O. compressus*, **ii**) *N. sammara* and **iii**) *M. violacea*. Rod banks are numbered as B_n_. Representative rod and cone outer segments are indicated by black and white arrows, respectively. **B**. Densities of different types of retinal cells in *O. compressus* (*n*=1), *N. sammara* (*n*=1) and *M. violacea* (*n*=1). PRL, photoreceptor layer; ONL, outer nuclear layer; INL, inner nuclear layer; GCL, ganglion cell layer; GC, ganglion cells. Scale bars: 25 μm (Ai), 50 μm (Aii and Aiii).

### Holocentrids have a higher temporal resolution compared to cardinalfish

Temporal resolution ERGs were conducted to determine the flicker fusion frequency (FFF; the point at which evenly spaced light pulses can no longer be distinguished as separate) in response to dim (4 lux) and bright (384 lux) stimuli at day (*n*=3) and night (*n*=5) (Fig. S2). Under all conditions, *N. sammara* attained the greatest FFF [mean ± s.e.m. at day and night, respectively: dim: 50±7.6 Hz and 33±3.7 Hz; bright: 70±2.9 Hz and 42.5±2.5 Hz; *p*<0.05 except for dim stimuli during the day which was not significant (n.s.)], followed by *M. violacea* (dim: 43.3±1.7 Hz and 20±0 Hz; bright: 57.5±2.5 Hz and 25±0 Hz) and then *O. compressus* (dim: 38.3±1.7 Hz and 17±2.5 Hz; bright: 41.7±1.7 Hz and 13±4.9 Hz) (Fig. 2; Fig. S3; Table S3). Furthermore, holocentrids had lower FFFs when exposed to the dim stimulus compared to the bright stimulus at each time point (*p*<0.05 for dim vs. bright stimulus during the day and dim vs. bright stimulus at night for both species; Table S3). However, the FFFs of *O. compressus* did not vary greatly with stimulus intensity. Finally, all species showed a trend towards lower FFFs at night compared to during the day, irrespective of stimulus intensity (*p*<0.0001 for day vs. night for bright stimulus and day vs. night for dim stimulus for all species; Table S3).

**Fig. 2.**
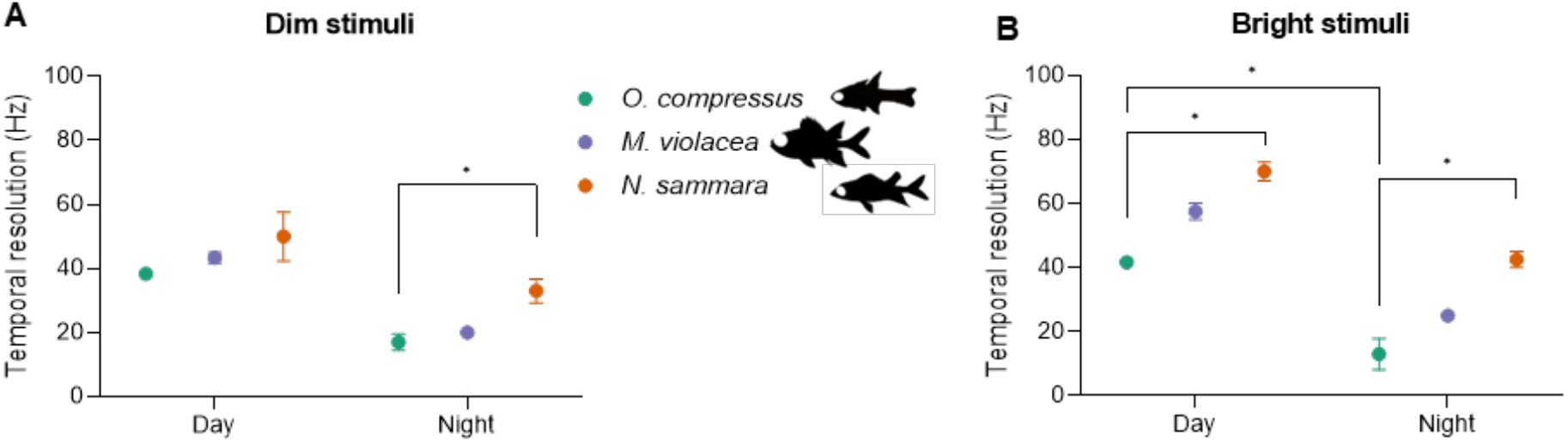
Temporal resolution electroretinography (ERG). ERG waveforms were obtained for a range of stimulus frequencies from 5 to 95 Hz. The temporal correlation of resultant waveforms with the stimulus were used to derive the maximal temporal resolution (*i*.*e*., flicker fusion frequency) elicited using either **A**) dim (4 lux) or **B**) bright (384 lux) stimuli in *O. compressus* (*n*=3 and 5 for day and night recordings, respectively), *N. sammara* (*n*=3 and 5 for day and night recordings, respectively) and *M. violacea* (*n*=3 and 5 for day and night recordings, respectively). Data represent mean ± s.e.m. Statistical significance (calculated from a Kruskal-Wallis with Dunn’s multiple comparisons test): *, *p*<0.05.

### Holocentrids have enhanced sensitivity compared to cardinalfish to both bright and dim light at night

Absolute sensitivity ERGs were recorded for *O. compressus* (*n*=5), *N. sammara* (*n*=4 and 5 for day and night recordings, respectively), and *M. violacea* (*n*=4) during the day and night (Fig. S2; Fig. S4). Firstly, *V/logI* curves were normalised to either V_max_ alone (for sensitivity of the entire eye; Fig. 3A) or V_max_ and eye size (for sensitivity per unit of retina; Fig. 3B). In all species, *V/logI* curves produced non-monotonic functions, with the amplitude of the b-wave representing the response post-synaptic to the photoreceptor, generally increasing with stimulus intensity until the maximal amplitude (V_max_) was reached, before subsequently decreasing due to bleaching. Notably, a subtle plateau occurred in the curves from *M. violacea* between stimulus intensities of ∼40 and 700 lux [equivalent to 1.6-2.8 log_10_(lux)], before continuing to increase until the response reached its peak. A closer examination of the ERG waveforms themselves revealed that, in all species, the speed of the visual response (*i*.*e*., time taken for the b-wave to reach its peak) became faster at higher intensities (Fig. S5). Additionally, the photoreceptor-derived component of the waveform (*i*.*e*., a-wave amplitude) also increased at higher intensities, very minimally in *O. compressus*, more substantially in *N. sammara* and greatly in *M. violacea* (Fig. S5).

**Fig. 3.**
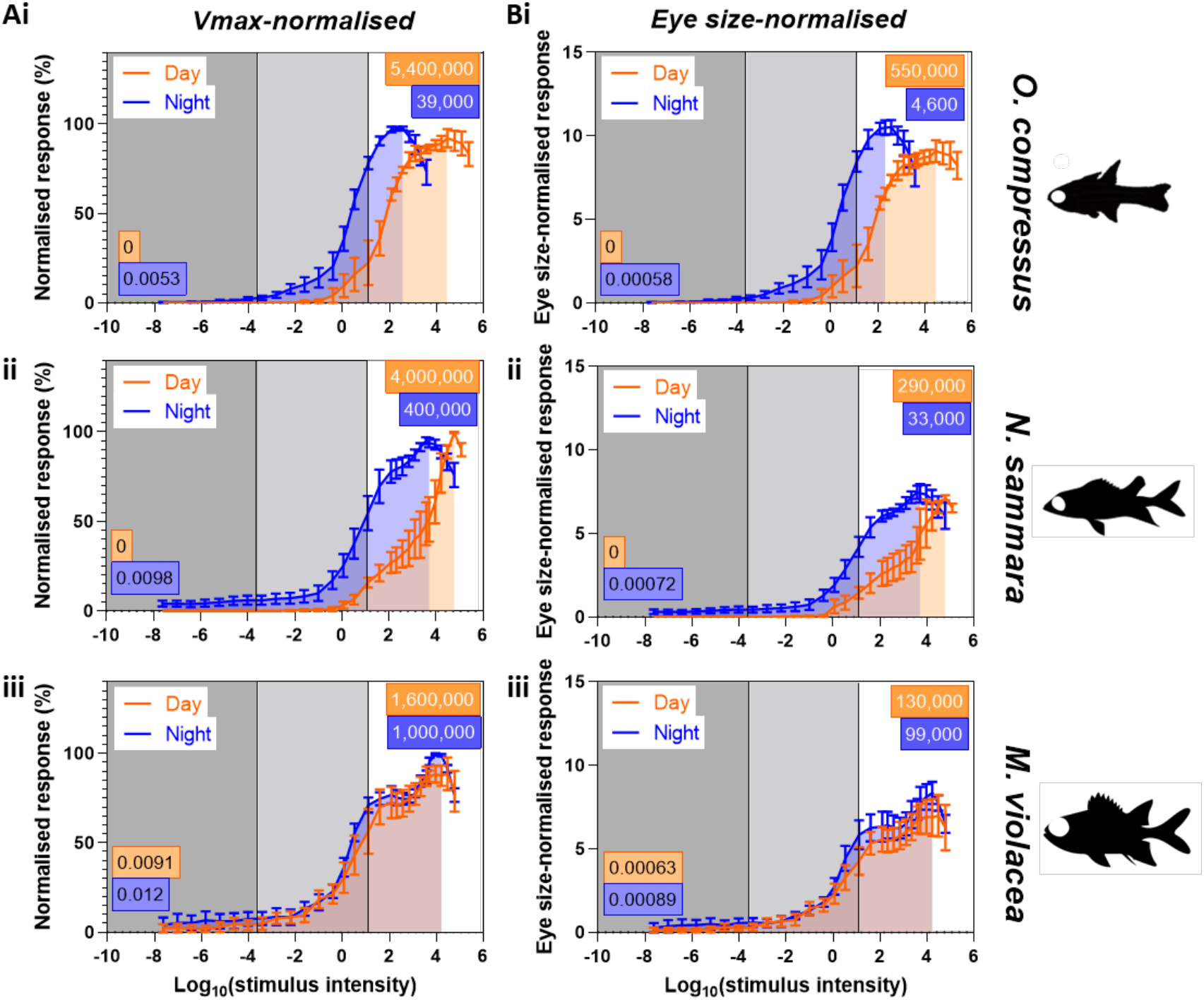
V/logI curves from absolute sensitivity electroretinography (ERG). ERG waveforms were obtained for a range of intensities from 2.4x10^−8^ to 240,000 lux [*i*.*e*., -7.6 to 5.4 log_10_(lux)] and the mean b-wave amplitude from each set of waveforms was plotted against the log_10_ of the stimulus intensity (in lux) normalised to either **A**) the maximal response (V_max_; response given as % of V_max_) or **B**) both V_max_ and eye size for **i**) *O. compressus* (*n*=5), **ii**) *N. sammara* (*n*=4 and 5 for day and night recordings, respectively), and **iii**) *M. violacea* (*n*=4) at day (orange) and night (blue). Each graph is divided into bright (white; >10 lux), intermediate (light grey; 0.002-10 lux) and dim (dark grey; <0.002 lux) intensities. Shaded regions under the line graphs represent the total area under the curve (AUC) from the lowest detectable response up to the maximal response. Values in the coloured boxes represent the rounded AUC values for dim (black text; bottom left) or bright intensities (white text; top right). Data are mean ± s.e.m.

There were notable differences in the *V/logI* curves between diel period and species. The *V/logI* curves were bright-shifted during the day compared to the night for *O. compressus* and *N. sammara*, but not *M. violacea*. Furthermore, when considering the same diel period, the *V/logI* curves differed between the three species, with the nature of these differences quantified using analyses of the area under the curve (AUC) within the intensity ranges of bright (>10 lux), dim (<0.002 lux) or overall (all intensities). Interspecific trends in the AUC values were the same irrespective of whether the data was normalised to V_max_ alone or V_max_ and eye size (Fig. 3; Table S4). Firstly, regardless of intensity category (*i*.*e*., overall, bright, or dim), *M. violacea* had the greatest AUCs during the night, followed by *N. sammara* and then *O. compressus* (Fig. 3, Table S4), indicating that the holocentrids were more sensitive to both bright and dim intensities during the night than *O. compressus*. At dim intensities during the day, *M. violacea* was the only species that had a calculable AUC, indicating that *M. violacea* was the only species sensitive to dim intensities during the day. Finally, for both overall and bright intensities, *O. compressus* had the greatest AUCs during the day, followed by *N. sammara* and then *M. violacea*, indicating that *O. compressus* was more sensitive to brighter intensities during the day than both holocentrids.

### Holocentrids had faster estimated retinal release kinetics compared to cardinalfish

The retinal release kinetics of each species’ rhodopsin protein were estimated using AA substitutions. The *O. compressus* RH1 possessed four AA substitutions known to alter retinal release rate, while those in *N. sammara* and *M. violacea* had six and seven AA substitutions, respectively (Table 1). These substitutions resulted in reduced estimated retinal release times for the rhodopsins of all three species when compared to wild-type rhodopsin. Estimations of the cumulative decrease in retinal release half-life were greatest in *M. violacea* (*t*_*1/2*_ difference of -6.6 min), followed by *N. sammara* (-5.3 min) and then *O. compressus* (-4.3 min). Therefore, the rhodopsins of both holocentrids had faster estimated retinal release kinetics than that of *O. compressus*.

**Table 1.** Amino acid substitutions in nocturnal reef fishes linked to retinal release kinetics. Different amino acid substitutions (AA) in teleosts that have been found to have little effect on spectral tuning but alter retinal release kinetics (53-55) were examined in *O. compressus, N. sammara* and *M. violacea*. Each candidate AA substitution is given in the first column, and the corresponding AA found in the study species is given for each site. Substituted sites in the study species are in bold. The influence on retinal release was defined as the difference in retinal release *t*_*1/2*_ (min) compared to wild-type rhodopsin.

| AA substitution | *O. compressus* | *N. sammara* | *M. violacea* | Influence on retinal release |
| --- | --- | --- | --- | --- |
| I209V | T | F | **V** | -1.3 |
| F213I | M | M | L | +1.5 |
| V266L | C | **L** | **L** | +1.7 |
| L290I | **I** | **I** | **I** | -1.3 |
| V286I | V | L | L | -0.9 |
| M123I | **I** | **I** | **I** | +4.9 |
| G124A | **A** | S | G | +1.8 |
| C165L | C | **L** | **L** | -0.9 |
| V189I | **I** | **I** | **I** | +2.5 |
| L59Q | L | L | L | -6.3 |
| Y74F | Y | Y | Y | -1.9 |
| N83D | **D** | **D** | **D** | -12.2 |
| **Cumulative change** | -4.3 | -5.3 | -6.6 |  |

## Discussion

Here, we investigated the retinal structure and visual function of nocturnal reef fishes with and without multibank retinas. Firstly, we confirmed that, at the morphological level, the three species investigated had visual systems that were well-adapted to their dim light environments (Fig. 1; Fig. S1; Table S2). In accordance with their nocturnal lifestyle (11, 12, 35), all three species had high rod densities and high rod:GC summation, and low cone and GC densities. Additionally, like other holocentrid species (12), *N. sammara* and *M. violacea* had multiple rod banks across the entire retina. Similar to other nocturnal reef fishes (11, 12), all three species also retained some degree of photopic adaptation, with cones organised into regional specialisations. However, the degree of scotopic and photopic adaptations varied between the three species with *N. sammara* and *M. violacea* showing greater adaptation for scotopic vision (*i*.*e*., higher rod densities and summation and multibank retinas) but inferior adaptation for photopic vision (*i*.*e*., lower cone densities) compared to *O. compressus*.

Secondly, this study examined temporal resolution (or speed) of vision in these fishes by determining the flicker fusion frequency (FFF) (Fig. 2; Fig. S3; Table S3). Temporal resolution is fundamentally determined by the integration time of photoreceptors, with cones displaying faster dynamics than rods (36). Thus, FFF is generally slower in conditions when rod responses dominate, such as in species with rod-dominated retinas (*e*.*g*., deep-sea fishes), at night and for lower stimulus intensities (22, 32). Consequently, the maximal FFF of deeper-dwelling and nocturnal fishes ranges from about 9 to 40 Hz, compared to the 40 to 100 Hz in shallow-dwelling diurnal fishes (32, 34, 37). Similar to findings in other fishes (38), the FFF of *O. compressus, N. sammara*, and *M. violacea* varied with diel period and stimulus intensity. All species had dim-stimulus night-time FFFs comparable to other nocturnal reef fishes, however, the peak FFF (*i*.*e*., elicited with bright stimuli during the day) only fit within the range for other nocturnal fishes for *O. compressus* (∼40 Hz) (33). FFF peaked at much higher values for both *N. sammara* (70 Hz) and *M. violacea* (∼60 Hz), falling within a range that is usually characteristic of diurnal fishes (33, 34). The fact that *O. compressus* had the highest cone and lowest rod densities but not the highest peak FFF implies that more complex neuronal mechanisms are at play in the holocentrids, likely due to the structure of the multibank retina. To our knowledge, the only other multibank representative whose temporal resolution has been assessed was that of a deep-sea fish (*Lepidocybium flavobrunneum*) which was slow-moving and had a much lower FFF [9 Hz; (22)]. It is possible that the higher temporal resolution in holocentrids may represent an adaptation for active life in shallow waters (39, 40).

Finally, we assessed luminous sensitivity (Fig. 3; Fig. S4; Table S4). In fishes, luminous sensitivity usually varies with diel period due to a dominance of cone- and rod-based responses at day and night, respectively (32, 36). Our findings revealed that *N. sammara* and *O. compressus* were no exception, showing higher bright-light sensitivity during the day but higher dim-light sensitivity during the night. However, the sensitivity of *M. violacea* was relatively constant. This indicates that *M. violacea* may only undergo a weak diel switch between photopic and scotopic systems. This is likely due to their lack of a well-developed photopic system to switch to, similar to some deep-sea fishes with pure rod retinas (41).

Luminous sensitivity also varies with retinal structure and ecology. For example, diurnal fish (with higher cone densities) have greater day-time bright-light sensitivity, while nocturnal fish (higher rod densities) have greater night-time dim-light sensitivity (31, 32). Similarly, this study found increasing dim-light sensitivity at night with increasing rod densities and rod banking. This supports the theory that the multibank retina enhances dim-light sensitivity. Conversely, our findings showed increasing bright-light sensitivity during the day with increasing cone densities, suggesting that the multibank retina has less involvement in photopic vision when cones can be used instead. Finally, increasing rod densities and banking (and decreasing cone densities) enhanced bright-light sensitivity at night when rod responses dominate. However, it is unlikely that holocentrids need to respond to any bright intensities at night. Instead, the rods in the multibank retina may be facilitating bright-light sensitivity simply when the use of cones is restricted (*e*.*g*., when the retina is rod-dominated in dim-light specialised species). Interestingly, this rod-based bright-light sensitivity seems to be masked by the higher bright-light sensitivity of the cones during the day, particularly in *N. sammara*. Notably, the potentially rod-based bright-light sensitivity of the holocentrids did not seem to grant them the same level of day-time bright-light sensitivity as a fish with higher cone densities. However, their level of sensitivity would likely still be sufficient to meet their day-time ecological demands, such as courtship and predator avoidance (42, 43). Hence, as previously proposed (44), this finding suggests that holocentrids use the different layers of rods to regenerate the visual response, permitting some rod-based vision under brighter intensities during the day.

Our study suggests that the rods in the holocentrid multibank retina can still function at brighter intensities. However, rhodopsin normally bleaches at high intensities. A key reason for this bleaching is the slower retinal release rate of rhodopsin compared to the cone opsins (45, 46). Amino acid-based estimations of retinal release in our study species revealed that the holocentrids may have accelerated retinal release kinetics compared to cardinalfishes and a wildtype reference rhodopsin, which would allow their rods to recover more rapidly post-bleaching (Table 1). Supporting a faster recovery rate in holocentrids, we also found higher temporal resolution at both day and night compared to *O. compressus* despite their less well-developed photopic visual systems. Furthermore, work in mice has shown that rods can recover and respond to bright intensities and that this is facilitated by more efficient post-bleaching regeneration (47, 48). Future work using *in vitro* regeneration experiments to test the retinal release kinetics of holocentrid RH1 visual pigments may be used to explain how their rods continue to function at brighter intensities.

Overall, our findings suggest a dual role for the multibank retina, where at dim intensities it functions to enhance photon capture while at bright intensities, it functions to regenerate the visual response, allowing the eye to function at both lower and higher intensities than a retina with a single rod bank. Enhanced visual functionality at both bright and dim light intensities aligns well with the ecology of holocentrids, since they are nocturnal foragers but are still somewhat active on the reef during the day (42). Our results strongly support one of the predominant theories on the function of the multibank retina (16). However, it still remains possible that the multibank retina also permits colour vision in dim light (20). This second theory may be investigated behaviourally in future work using accessible, easy-to-maintain species with multibank retinas, such as the holocentrids.

## Materials and Methods

### Animal collection and ethics

Details of all animals are given in Table S1. Adult fish were collected from the Great Barrier Reef around Lizard Island, Australia or sourced from a supplier, Cairns Marine, which also collects from the northern Great Barrier Reef. All collections and procedures were conducted under a Great Barrier Reef Marine Park Permit (G17/38160.1), a Queensland General Fisheries Permit (180731), and a University of Queensland’s Animal Ethics Permit (QBI 304/16). Following euthanasia, all animals were photographed with a scale reference to quantify body length and eye diameter. Eyes were dissected and the eye cup preserved in RNAlater or paraformaldehyde [PFA; 4% (w/v) PFA in 0.01M phosphate-buffered saline (PBS), pH 7.4] depending on the analyses.

### Histology

Five retinal regions (dorsal, ventral, central, nasal and temporal) were dissected, processed and sectioned from PFA-fixed eyes as described in (12). The densities of key retinal cell types (*i*.*e*., cones, rods, INL cells and GC) per 0.01 mm^2^ of retina were estimated from sections using Fiji v1.53c (49) as described elsewhere [SI Appendix; (29)]. Densities were corrected for cell size using Abercrombie’s correction (50) (Fig. 1).

### Electroretinography (ERG)

Corneal ERG recordings were conducted *in vivo* on whole, intact eyes to assess visual function using methods similar to those described in (33). Fish were acclimatised to the recording chamber for 30 min, anaesthetised with 0.2 mL clove oil/litre seawater, immobilised with an intramuscular injection of 8.5 mg/kg gallamine triethiodide and ventilated with oxygenated seawater (Fig. S2). After ≥40 min of dark adaptation, light stimuli were delivered to the eye using a custom-built, calibrated, broad-spectrum light source controlled via a PowerLab 4/26 DAQ module. Visual responses were detected through silver wire electrodes placed on the surface of the eye, amplified via a DP-103 amplifier and acquired in LabChart 8 v8.1.16. The system was grounded to the water of the recording chamber. Recordings were conducted at 28 ± 1°C at both day and night to control for any effects of temperature and circadian rhythm, respectively. Recordings were performed at the Lizard Island Research Station (LIRS) or the Queensland Brain Institute (QBI). Additional recordings were taken at both sites to compare results between the recording locations (Fig. S6).

### Temporal resolution ERGs

The temporal resolution of vision was assessed using flicker fusion frequency (FFF) ERGs. FFF is the point at which evenly spaced light pulses can no longer be distinguished as separate. Dark-adapted FFF ERGs were recorded by increasing the frequency of white light stimuli of constant intensity from 5 Hz to 95 Hz at increments of 5 Hz. Light pulses were 10 ms in duration and were repeated 30 times. Recordings were conducted for bright (384 lux) and dim (4 lux) stimuli (Fig. 2). The FFF threshold was determined either through visual inspection (at lower frequencies, <65 Hz) or by using the power spectrum to differentiate the signal and noise (at higher frequencies, ≥65 Hz) [SI Appendix; (34, 51)]. Statistics and graphs throughout the study were generated in GraphPad Prism v9.0.0.

### Absolute sensitivity ERGs

The absolute (luminous) sensitivity of vision was determined using *V/logI* curves, which plot the normalised amplitude of the response, *V* (Fig. S2), against the log of the intensity (*I*). These ERGs were recorded by increasing the intensity of a white light from 2.4x10^−8^ to 240,000 lux [*i*.*e*., -7.6 to 5.4 log_10_(lux)] in 0.3-0.6 log unit steps. Light stimuli were 100 ms pulses presented at 0.1 – 0.4 Hz (SI Appendix) and were repeated ten times for each intensity. The mean response amplitudes were normalised to the maximal response (V_max_) and plotted against stimulus intensity to obtain the *V/logI* curve (33, 52). The area under the curve (AUC) was calculated as a proxy for the magnitude and breadth of the visual responses. AUC was calculated for either all intensities, dim intensities (<0.002 lux) or bright intensities (>10 lux) for each species (Fig. 3). To isolate the effect of the multibank retina, the V_max_-normalised responses were also normalised to eye size (to obtain responses per unit of retina) and analysed again as described above. To further understand how the visual response changed with intensity, representative ERG waveforms were analysed to obtain: 1) the time from stimulus presentation to the peak of the signal generated post-synaptic to the photoreceptors (*i*.*e*., time to b-wave peak; ms) and 2) the amplitude of the photoreceptor-derived peak (*i*.*e*, a-wave amplitude; mV). These values were obtained for dim (0.4 lux), moderate (125 lux) and bright (2165 lux for *O. compressus* and 5160 lux for *N. sammara* and *M. violacea*) stimuli, which matched the base, peak and decline of the *V/logI* curves, respectively.

### Estimations of retinal release kinetics

Amino acid substitution sites involved in retinal release kinetics were used to estimate the retinal release time of the rhodopsin protein in each species (Table 1). Firstly, 12 candidate amino acid (AA) substitution sites were identified from the literature (53-55). Notably, retinal release effect has not been characterised for all positively selected non-spectral substitutions in the literature (*e*.*g*., T97S in *N. sammara* and F116S and A164G in *M. violacea*) and that any substitutions that also affected spectral sensitivity were excluded from these analyses. Next, the rhodopsin coding sequences for *O. compressus* (MH979489.1), *N. sammara* (MW219675.1) and *M. violacea* (MW219672.1) (11, 12) were downloaded from GenBank (https://www.ncbi.nlm.nih.gov/genbank/) and translated to protein sequences. These were manually inspected for AA substitutions at each of the 12 candidate sites in Geneious Prime v2021.1.1. Identified substitutions were used to estimate the cumulative change in retinal release, calculated as the difference in retinal release half-life (*t*_*1/2*_; min) compared to wild-type zebrafish (53), bovine (55) or catfish (56) rhodopsin, depending on the study (Table 1).

## Acknowledgements

We acknowledge the Dingaal, Ngurrumungu and Thanhil peoples as traditional owners of the lands and waters of the Lizard Island region from where specimens were collected. We also acknowledge the traditional owners of the land on which the University of Queensland is situated, the Turrbal/Jagera people. We would like to thank Cairns Marine for supplying animals and the staff at Lizard Island Research Station, Anne Hoggett and Lyle Vail, for support during field work. We thank Robert Sullivan from the Queensland Brain Institute (QBI) Histology Facility, Richard Webb and Robyn Chapman Webb from the Centre for Microscopy and Microanalysis (CMM) and Rumelo Amor from the QBI Advanced Microscopy facility for technical support and advice. Finally, we thank Dr Martin Luehrmann for invaluable guidance and discussions about the findings.

## Data Availability

All study data are included in the article and/or SI Appendix.

