## Supplementary Information for "Multiple rod layers increase the speed and sensitivity of vision in nocturnal reef fishes"

### Expanded Materials and Methods

**Animal collection and ethics.** Three species of nocturnal coral reef fishes, *Ostorhinchus compressus*, *Neoniphon sammara* and *Myripristis violacea*, were investigated in this study. Details of all animals used in this study are given in Table S1. All *N. sammara* and *M. violacea* and some *O. compressus* were collected with clove oil and hand nets on the Great Barrier Reef around Lizard Island, Australia under a Great Barrier Reef Marine Park Permit (G17/38160.1) and Queensland General Fisheries Permit (180731). Most *O. compressus* were sourced from a supplier that collects from the northern Great Barrier Reef, Cairns Marine (https://www.cairnsmarine.com/). After being used in electroretinography experiments, all individuals were first anaesthetized by immersion in a solution of 0.2ml clove oil/L seawater until respiration and response to light and touch had ceased and were then euthanized by swift decapitation. Following euthanasia, all individuals were photographed with a scale reference (to quantify body length and eye diameter) and eyes were immediately enucleated, the cornea and lens removed, and the eye cup preserved in either RNAlater (Sigma-Aldrich) or 4% paraformaldehyde [PFA; 4% (w/v) PFA in 0.01M phosphate-buffered saline (PBS), pH 7.4] depending on the analysis (see below for details). All procedures were approved by the University of Queensland Animal Ethics Committee (QBI 304/16).

**Histological analyses.** One PFA-fixed eye from *N. sammara*, *M. violacea* and *O. compressus* was used for histological analysis. The retinas were dissected out of the eye cup and a square of retina was dissected from each of five key retinal regions (dorsal, ventral, central, nasal and temporal) to account for intraretinal variability (1) (Figure S1). Each piece of retina was post-fixed in 2.5% glutaraldehyde and 2% osmium tetroxide, progressively dehydrated in increasing concentrations of ethanol and infiltrated with Epon resin (ProSci Tech) in a BioWave Pro tissue processor. Resin samples were then polymerized at 60°C for 48 h. 1 μm-thick radial sections were cut on a Leica ultramicrotome (Ultracut UC6), de-plasticised in 11% sodium ethoxide, rehydrated and stained with a solution of 0.5% toluidine blue and 0.5% borax. Sections were captured using an Axio upright brightfield microscope (Zeiss, Imager Z1) with a 63X objective (oil, 1.4 numerical aperture, 0.19 mm working distance, 0.102 μm/pixel).

Cell densities were estimated from sections using the image analysis software, Fiji v1.53c (2). Briefly, each image was cropped into a retinal counting strip of 250 μm (horizontal length) for lower-density cell types (*i.e.,* cone and GCs) or 40 μm for the higher-density cell types (*i.e.,* ONL and INL cells). These counting strips were optimised for each cell type as described elsewhere (3). The number of cone outer segments (OS), outer nuclear layer (ONL) nuclei, inner nuclear layer (INL) nuclei and ganglion cell layer (GCL) nuclei were counted for three sections per sample using the cell counter plugin in Fiji. Cell counts were averaged and multiplied by a factor of 40 or 250 (depending on counting frame width) and divided by section thickness to obtain the density of each retinal cell type per 0.01 $mm^2$ of retina. Densities were corrected for cell size using Abercrombie's correction (4). Finally, rod densities were calculated as the difference between the number of ONL nuclei and the number of cone OS.

**Electroretinography (ERG).** Corneal ERGs were conducted *in vivo* on whole, intact eyes to determine the absolute (luminous) sensitivity and the temporal resolution of vision in *N. sammara*, *M. violacea* and *O. compressus*. ERG recordings capture the summed electrical potentials of the whole retina and as such represent a holistic approach to assessing visual responses, also accounting for the effects of ocular filtering and intraretinal processing (5). Furthermore, this technique permits inter-species comparisons (6-8). Methods were similar to those described in (6), with minor modifications. Firstly, each fish was acclimatised to the temperature of the recording chamber water for 30 min, anaesthetised using 0.2 mL clove oil/litre seawater and prepared for ERG recordings using methods approved by the University of Queensland Animal Ethics Committee (QBI 304/16). Fish were immobilised with an intramuscular injection of 8.5 mg/kg gallamine triethiodide (Flaxedil; Sigma) and ventilated with oxygenated seawater (flow rate: 1-2 L/min) (Figure S2).

Following at least 40 min of dark adaptation, a broad-spectrum (white) LED-based light source (built in-house with controllable voltage and a mirror-focussed beam) was placed 10 cm above the eye of the fish to deliver light stimuli. The irradiance of the light was calibrated with a Jaz spectrometer (Ocean Optics). The light was connected to a PowerLab 4/26 DAQ module (ADinstruments) which allowed control of the intensity, frequency, and duration of light stimuli via the LabChart 8 software v8.1.16 (ADinstruments). Visual signals were detected through Teflon-coated, chlorided silver wire electrodes (0.5mm; Ag-AgCl2) placed on the corneal surface of the eye (active electrode) and the fatty tissue within the orbit (reference electrode) (Figure S2). Signals were recorded at a sampling frequency of 4 kHz and then amplified and pre-filtered using a DP-103 amplifier (Warner Instruments) set to 1000x gain and 1 Hz high pass and 1 kHz low pass filters. Amplified signals were further filtered in LabChart using a notch filter at 50Hz to remove electrical noise. The system was grounded to the water of the recording chamber.

Seawater in the recording chamber was maintained at a constant temperature of 28 ± 1˚C to control for any effects of temperature on the visual response. Recordings were conducted at day and night to evaluate any effects of circadian rhythm. All recordings were conducted between February and December 2020. Fish collected around Lizard Island were maintained in outdoor aquaria onsite, while fish from Cairns Marine were housed in indoor aquaria at The University of Queensland. All fish were maintained in a normal photoperiod of 12 h of light and 12 h of dark for indoor aquaria, and 12 h 40 min of light and 11 h 20 min of dark for outdoor aquaria. ERGs on *M. violacea* and *N. sammara* were conducted at the Lizard Island Research Station (LIRS) and recordings on *O. compressus* were conducted at the Queensland Brain Institute (QBI). Additional recordings on *O. compressus* were conducted at LIRS to confirm that the recording location did not affect the visual response and that results between species were comparable (Figure S6). All ERG data were analysed in LabChart and GraphPad Prism software v9.0.0 (www.graphpad.com).

**Temporal resolution ERGs.** The temporal resolution of vision was assessed using flicker fusion frequency (FFF) ERGs. FFF is the frequency at which evenly spaced flashes of light are no longer sufficiently temporally separated for the eye to resolve. This translates on the recordings as ERG waveforms that no longer follow the flickering of the light source. FFF ERGs were recorded in the dark-adapted state as the frequency of a white light pulse of constant intensity was increased from 5 Hz to 95 Hz at increments of 5 Hz. Light stimuli were 10 ms in duration and were repeated 30 times at each frequency. Recordings were conducted for bright (384 lux) and dim (4 lux) stimuli. The number of individuals tested per species and condition was 3 and 5 for the day and night recordings, respectively.

The FFF threshold was determined using two methods. The first method involved visual inspection of ERG waveforms to determine if they followed the flickering stimuli (Figure S2). This method was only possible at lower frequencies and was therefore, only used for stimuli less than 65 Hz. A second, standardised method was employed for frequencies above 65 Hz (9, 10). Briefly, the FFF threshold was determined by analysing the power spectrum of averaged responses. The power at the stimulus frequency (signal) was compared to that at a neighbouring, non-stimulus frequency (noise). The FFF threshold was defined as the highest frequency at which the signal was five times stronger than the noise.

**Absolute sensitivity ERGs.** The absolute (luminous) sensitivity of vision was determined using intensity-response (*V/logI*) curves. A *V/logI* curve plots the normalised amplitude of the visual response (measured as the amplitude of the b-wave; Figure S2) at the log of a given intensity (*I*). Stimuli were presented across a range of intensities from $2.4 \times 10^{-8}$ to 240,000 lux [*i.e.,* -7.6 to 5.4 $\log_{10}$(lux)] in 0.3-0.6 log unit steps. This range covered undetectable responses through to the maximal response and into the bleached state in all animals. At each intensity step, the ERG response to ten repeated 100 ms flashes was recorded with 2.4 – 9.9 sec rest intervals (*i.e.,* presented at a frequency of 0.1 – 0.4 Hz), depending on the intensity step. The stimulus frequency at each intensity was determined experimentally prior to the trials by assessing the amplitude of the b-wave with different recovery times and using the shortest interval (*i.e.,* highest frequency) that restored the maximal amplitude. Different intensities were exacted by varying the voltage of the LED and placing neutral density (ND) filters (0.3 and 0.6; LEE Filters) in the light path. The mean b-wave peak amplitude was calculated at each intensity and this was normalised to the maximal response ($V_{max}$) and plotted against stimulus intensity to obtain the *V/logI* curve (6, 11). Recordings were conducted for *N. sammara* (*n*=4 during the day, *n*=5 at night), *M. violacea* (*n*=4) and *O. compressus* (*n*=5).

As luminous sensitivity encompasses both the magnitude and breadth of the visual response, the area under the *V/logI* curve (AUC) was used as a combined statistic conveying both mean normalised response amplitude and the breadth of intensities that the fish was responsive to. Thus, the AUC was calculated separately at either all intensities, dim intensities only (taken as <0.002 lux) or bright intensities only (>10 lux) for each species. To further understand how the visual response might be changing at different intensities, representative ERGs were used to obtain: 1) the time between stimulus delivery and the peak of the ERG component post-synaptic to the photoreceptors (*i.e.,* time to b-wave peak; ms) and 2) the amplitude of the photoreceptor-derived component of the ERG (*i.e,* a-wave amplitude; mV). These parameters were calculated for dim (0.4 lux), moderate (125 lux) and bright (2165 lux for *O. compressus* and 5160 lux for *N. sammara* and *M. violacea*) stimulus intensities. These specific intensities were chosen from *V/logI* curves as they occur at the base of the *V/logI* curve, just prior to reaching $V_{max}$ (*i.e.,* just prior to the curve peak) and just after $V_{max}$ is reached, respectively. Finally, to isolate the impact of a multibank retina on sensitivity, the $V_{max}$-normalised responses were also normalised to eye size and analysed again as described above.

**Estimations of retinal release kinetics.** Amino acid substitution sites involved in retinal release kinetics were used to estimate retinal release time of the rhodopsin protein expressed in each species used in this study. Firstly, amino acid (AA) substitutions, known to have little effect on spectral tuning but to alter retinal release kinetics in teleosts, were identified from the literature. This search yielded 12 candidate AA sites with experimentally tested effects on retinal release time (12-14). Published rhodopsin coding sequences for *O. compressus* (MH979489.1), *N. sammara* (MW219675.1) and *M. violacea* (MW219672.1) (1, 15) were then downloaded from GenBank (https://www.ncbi.nlm.nih.gov/genbank/). These coding sequences were then translated to obtain protein sequences and manually inspected for AA substitutions at each of the 12 candidate sites in Geneious Prime v2021.1.1 (Biomatters Ltd). Finally, the characterised effects of these substitutions were used to estimate the cumulative change in retinal release, and this was calculated as the difference in retinal release $t_{1/2}$ (min) compared wild-type zebrafish (12), bovine (14) or catfish (16) rhodopsin, depending on the study.

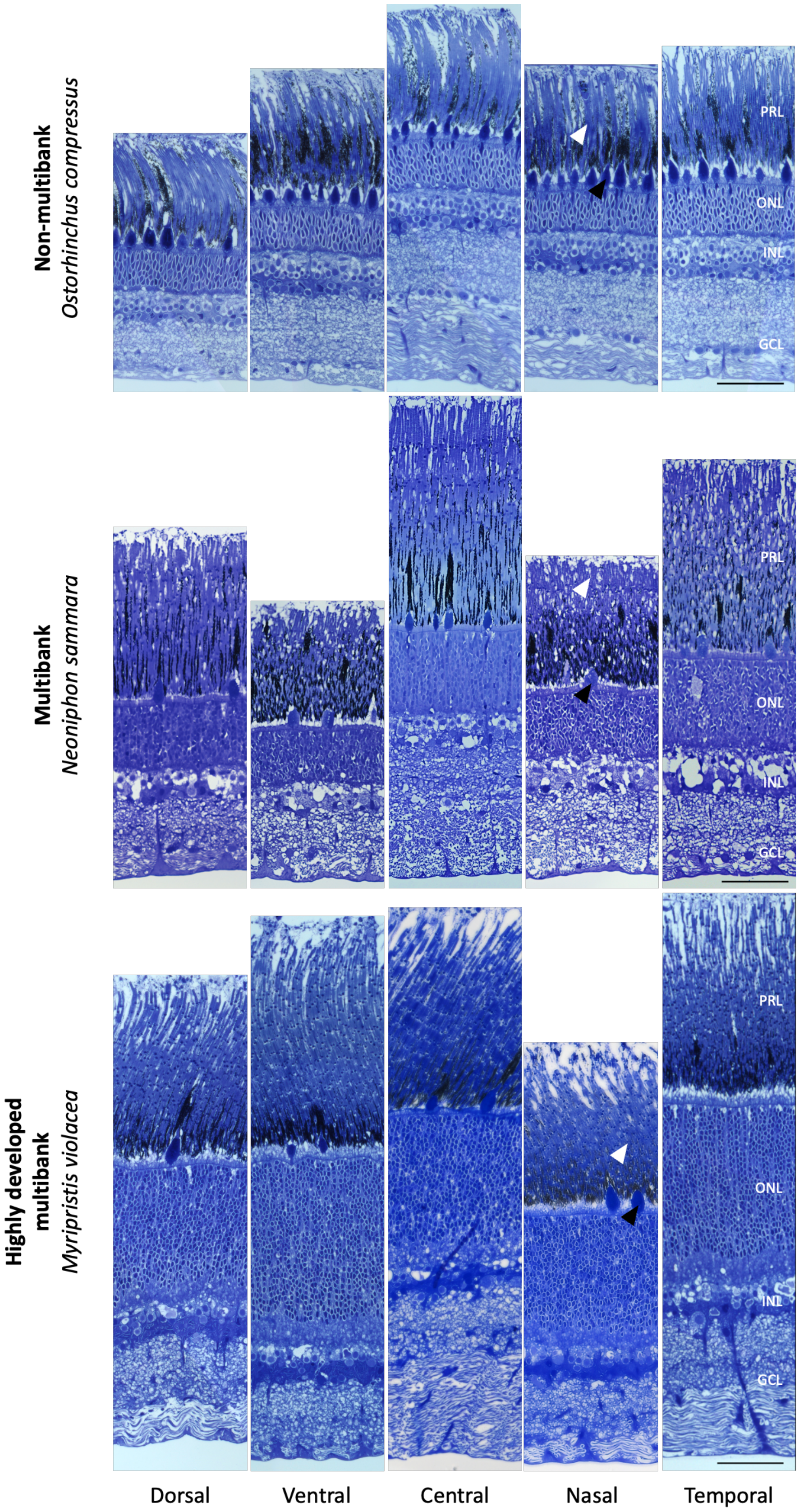


*Fig. S1. Retinal structure in O. compressus, N. sammara and M. violacea.* Representative radial retinal sections from five key retinal regions showing variation in rod banking between species and retinal regions. Representative rod and cone outer segments are labelled with white and black arrows, respectively. PRL, photoreceptor layer; ONL, outer nuclear layer; INL, inner nuclear layer; GCL, ganglion cell layer; GC, ganglion cells. Scale bars: 50 µm.

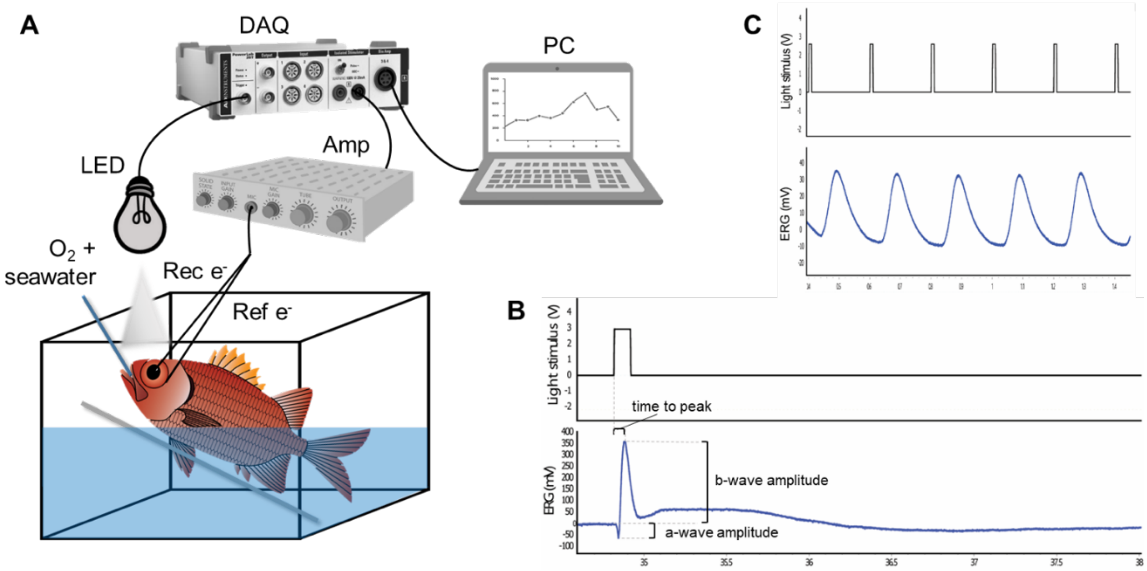


*Fig. S2. ERG recording setup.* **A.** Graphical representation of recording setup. **B.** Representative waveform from a luminous sensitivity ERG illustrating the amplitude of the b-wave (used to construct the *V/logI* curves) and the a-wave (used to gain an insight into the strength of the photoreceptor response) and the time to peak (*i.e.,* time between stimulus delivery and the peak of the b-wave). **C.** Representative waveforms from a temporal resolution ERG at a low stimulus frequency which the fish can resolve, illustrating how the visual response follows the stimulus. Rec e-, recording electrode; Ref e-, reference electrode; LED, light-emitting diode; DAQ, data acquisition card; Amp, amplifier; PC, computer.

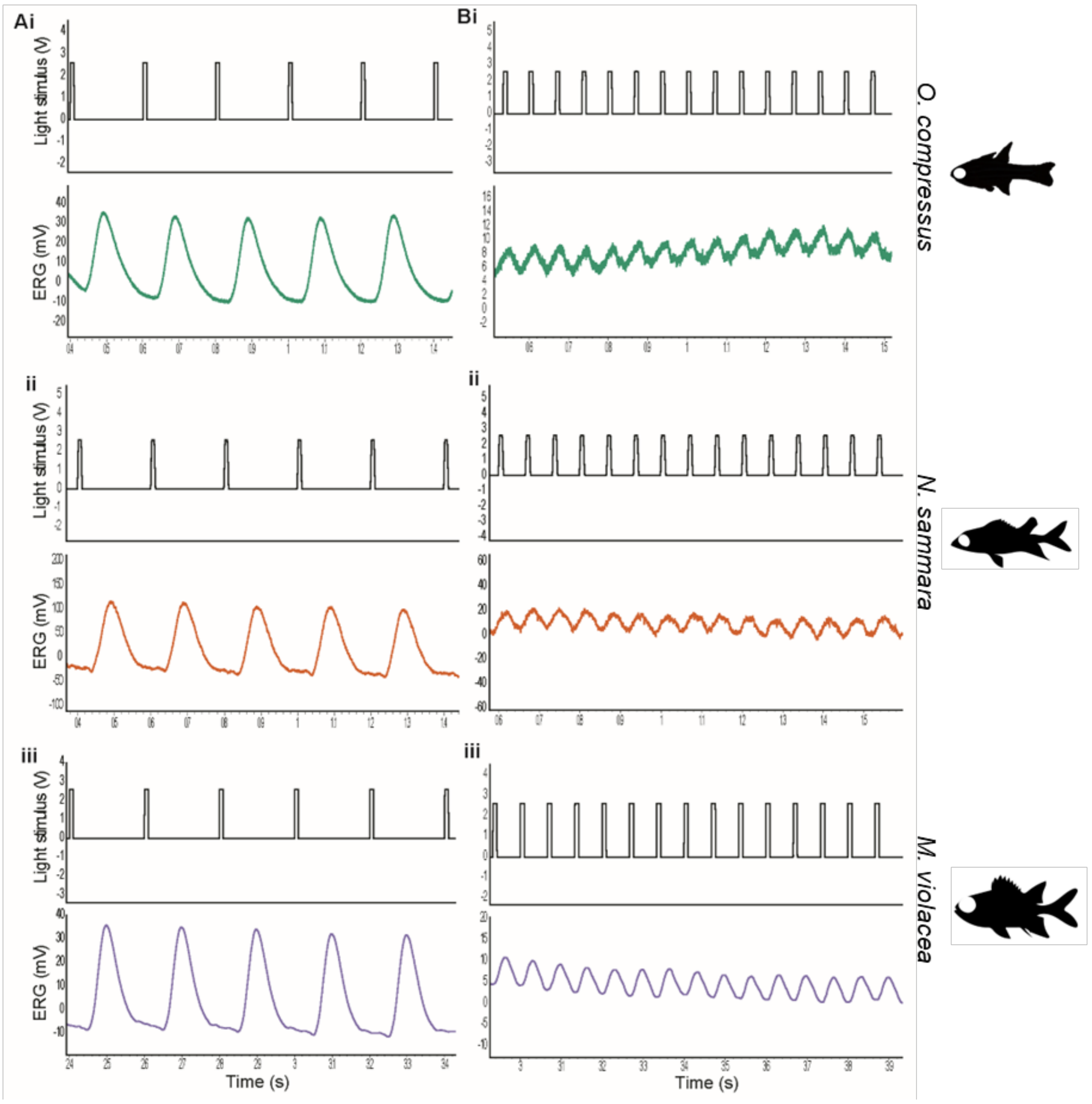


*Fig. S3. Temporal resolution electroretinography (ERG) waveforms.* Representative waveforms from temporal resolution ERGs for **i**) *O. compressus*, **ii**) *N. sammara,* and **iii**) *M. violacea* exposed to a 40-lux stimulus flickering at **A**) 5 Hz or **B**) 15 Hz.

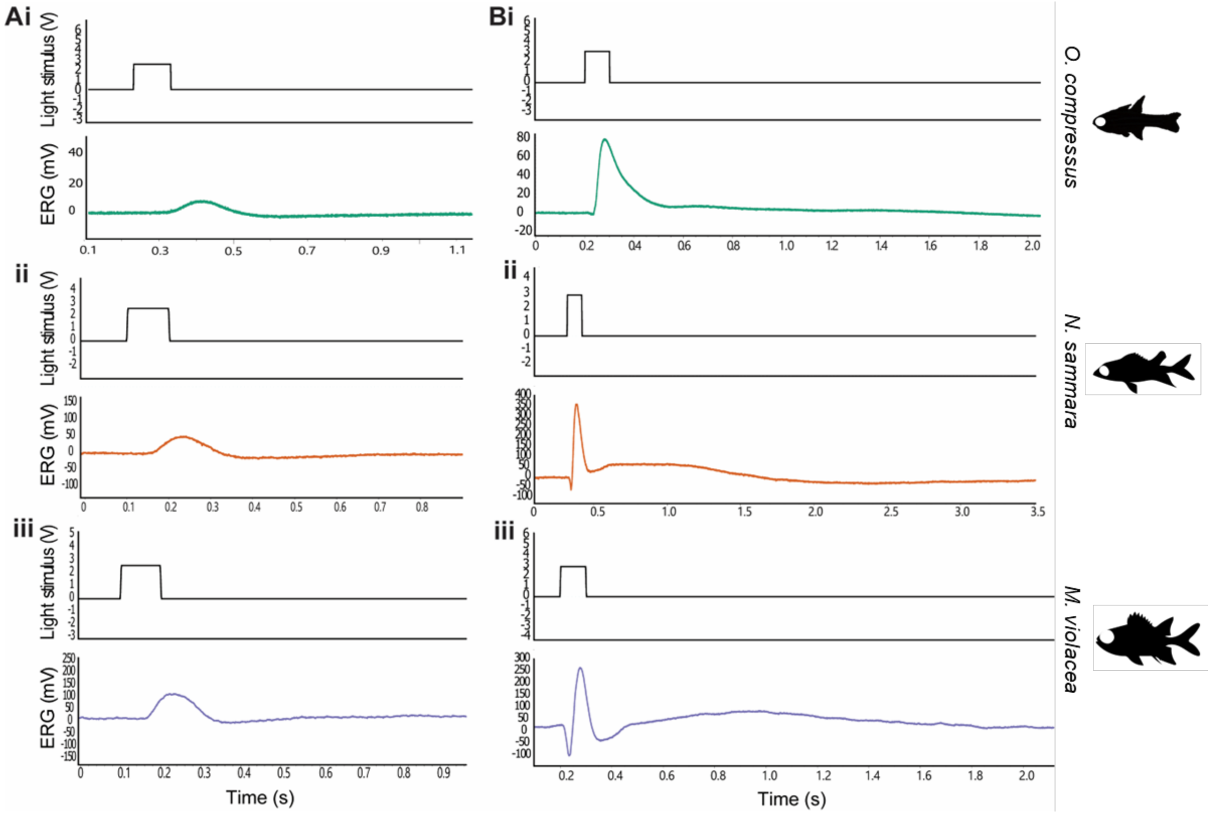


*Fig. S4. Absolute sensitivity electroretinography (ERG) waveforms.* Representative waveforms from absolute sensitivity ERGs for **i**) *O. compressus*, **ii**) *N. sammara*, and **iii**) *M. violacea* exposed to **A**) dim [0.0015625 lux, *i.e.,* -2.8 $\log_{10}$(lux)] or **B**) bright [375 lux, *i.e.,* 2.6 $\log_{10}$(lux)] light stimuli.

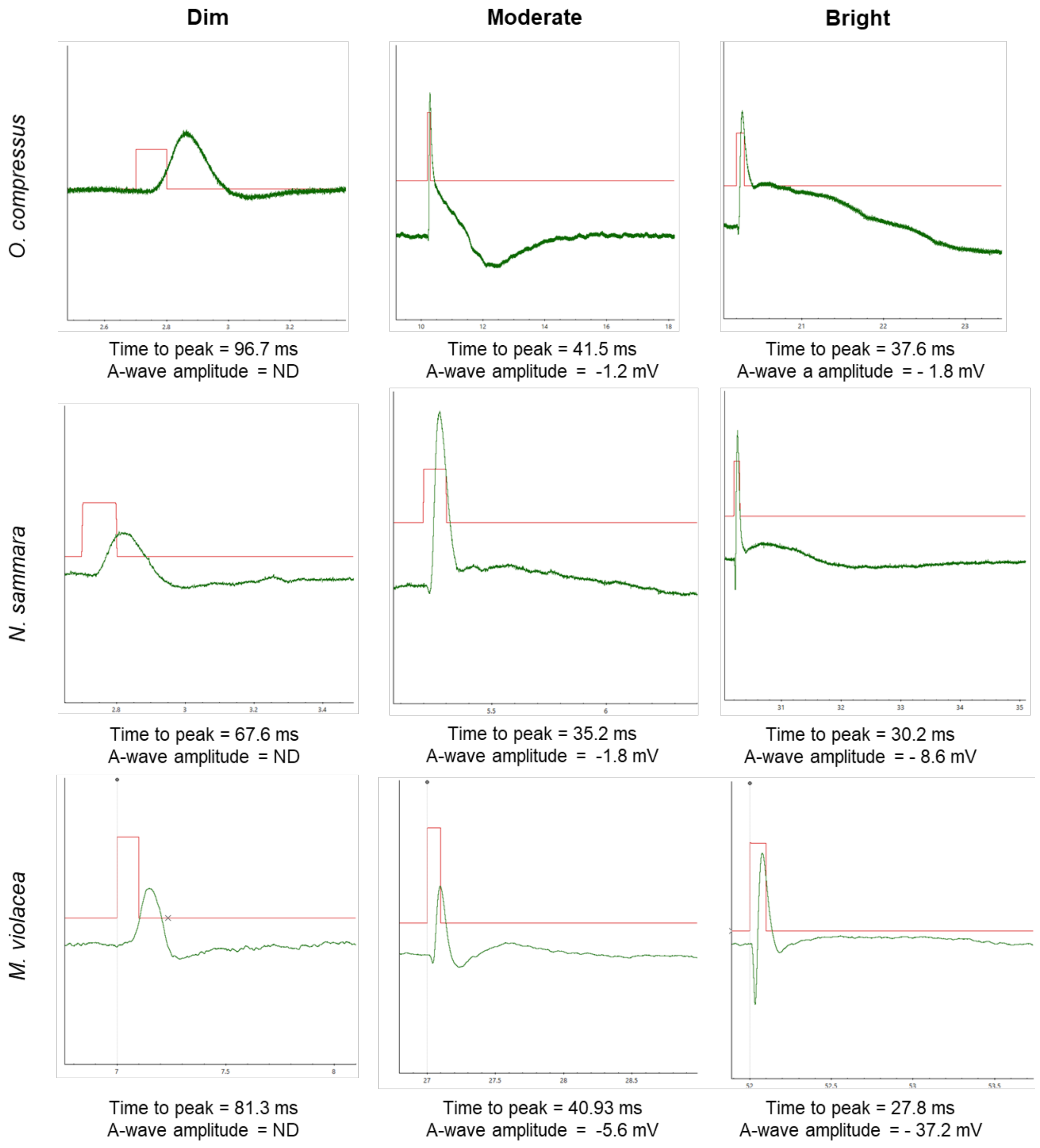


*Fig. S5. Time to peak and a-wave amplitude in representative luminous sensitivity ERGs from* O. compressus*,* N. sammara *and* M. violacea*.* Representative responses from luminous sensitivity ERGs from the three species with time to peak (ms) and a-wave amplitude (mV) given below. Responses are to dim (0.4 lux), moderate (125 lux) and bright (2165 and 5160 lux for cardinalfish and holocentrids, respectively) stimuli as indicated. These stimulus intensities were chosen from *V/logI* curves as they represent intensities at the base of the curve, just prior to reaching $V_{max}$ (*i.e.,* just prior to the curve peak) and just after $V_{max}$ is reached, respectively.

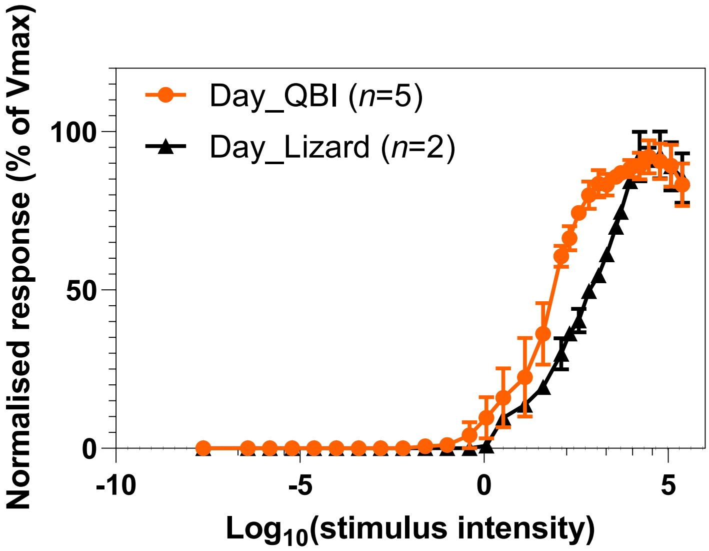


*Fig. S6. V/logI curves for O. compressus from two recording locations.* Comparison of *V/logI* curves (mean ± s.e.m.) derived from ERG recordings on *O. compressus* conducted at the Queensland Brain Institute (QBI) or the Lizard Island Research Station (LIRS). Data collected at QBI was used for comparison to other two species and data from LIRS was used to confirm that results were comparable between the two locations and that changing recording location did not significantly affect the visual response. In particular, note that the lower threshold of the response and the maximal response occurred at similar stimulus intensities in both locations.

*Table S1. Details of animals used in study.* All animals were collected from the northern Great Barrier Reef by either Cairns Marine (CM) or by the authors from the reefs surrounding Lizard Island Research Station (LI). Standard length (SL) and eye diameter are both given in mm. The time of ERG recording for each fish is also given.

| **Species** | **Recording time** | **SL (mm)** | **Eye diameter (mm)** | **Collection** |
|---|---|---|---|---|
| *Ostorhinchus compressus* | Day | 76.3 | 9.6 | CM |
| | | 93.7 | 11.2 | CM |
| | | 95.7 | 11.9 | CM |
| | | 81.8 | 9.8 | CM |
| | | 78.0 | 8.6 | CM |
| | | 77.6 | 10.5 | LI |
| | | 60.9 | 8.8 | LI |
| | Night | 86.5 | 9.1 | CM |
| | | 91.9 | 10.1 | CM |
| | | 92.2 | 10.3 | CM |
| | | 80.0 | 8.4 | CM |
| | | 79.0 | 8.8 | CM |
| *Myripristis violacea* | Day | 126.7 | 15.3 | LI |
| | | 76.4 | 10 | LI |
| | | 100.5 | 12.4 | LI |
| | | 109 | 15.7 | LI |
| | Night | 123.1 | 12.8 | LI |
| | | 114.0 | 11.7 | LI |
| | | 81.9 | 9.9 | LI |
| | | 99.4 | 14.4 | LI |
| *Neoniphon sammara* | Day | 104.4 | 9.2 | LI |
| | | 134.1 | 13.7 | LI |
| | | 132.8 | 14.6 | LI |
| | | 125.9 | 13.8 | LI |
| | | 122.0 | 14.6 | LI |
| | Night | 119.4 | 11.7 | LI |
| | | 116.5 | 14.7 | LI |
| | | 120.0 | 14.4 | LI |
| | | 101.4 | 12 | LI |
| | | 113.5 | 11.6 | LI |

*Table S2. Abercrombie-corrected retinal cell densities for* O. compressus*,* N. sammara *and* M. violacea*.* Densities of rods, cones, inner nuclear layer (INL) cells and ganglion cells (GC) and ratio of rods to GC in five key retinal regions: dorsal, ventral, central, nasal and temporal. Data are given in cells/0.01mm$^2$ (*n*=1).

| | **Species** | **Dorsal** | **Ventral** | **Central** | **Nasal** | **Temporal** |
|---|---|---|---|---|---|---|
| **Rods** | *O. compressus* | 3231 | 2057 | 3271 | 1740 | 3545 |
| | *N. sammara* | 8684 | 9662 | 10569 | 10265 | 12403 |
| | *M. violacea* | 17011 | 21196 | 18884 | 16569 | 18413 |
| **Cones** | *O. compressus* | 58.1 | 61.7 | 69.4 | 48.5 | 72.8 |
| | *N. sammara* | 18.4 | 22.3 | 38.8 | 26.2 | 49.4 |
| | *M. violacea* | 12.7 | 10.3 | 19.5 | 18.1 | 16.2 |
| **INL** | *O. compressus* | 761 | 930 | 971 | 530 | 1108 |
| | *N. sammara* | 659 | 474 | 676 | 629 | 789 |
| | *M. violacea* | 538 | 638 | 509 | 599 | 516 |
| **GC** | *O. compressus* | 41.4 | 59.1 | 89.3 | 50.3 | 97.5 |
| | *N. sammara* | 39.8 | 29.1 | 52.8 | 36.4 | 71 |
| | *M. violacea* | 12.1 | 12.8 | 29.2 | 23.3 | 22.9 |
| **Rods:GC** | *O. compressus* | 78.1 | 34.8 | 36.6 | 34.6 | 36.4 |
| | *N. sammara* | 218.3 | 332.6 | 200.2 | 282.3 | 174.8 |
| | *M. violacea* | 1403.4 | 1651.5 | 646.4 | 711.7 | 803.5 |

*Table S3. Temporal resolution ERGs.* Data are FFF calculated from temporal resolution ERGs in *O. compressus* (*n* = 3 and 5 for day and night recordings, respectively), *N. sammara* (*n* = 3 and 5 for day and night recordings, respectively) and *M. violacea* (*n* = 3 and 5 for day and night recordings, respectively) with dim (4 lux) or bright (384 lux) stimulus intensities. Data are mean ± SEM.

| **Stimulus intensity** | **Time-point** | ***O. compressus*** | | ***N. sammara*** | | ***M. violacea*** | |
|---|---|---|---|---|---|---|---|
| | | Mean | SEM | Mean | SEM | Mean | SEM |
| *Dim* | *Day* | 38.3 | 1.7 | 50.0 | 7.6 | 43.3 | 1.7 |
| | *Night* | 17.0 | 2.5 | 33.0 | 3.7 | 20.0 | 0.0 |
| *Bright* | *Day* | 41.7 | 1.7 | 70.0 | 2.9 | 57.5 | 2.5 |
| | *Night* | 13.0 | 4.9 | 42.5 | 2.5 | 25.0 | 0.0 |

*Table S4. Absolute sensitivity ERGs.* Data are areas under the curve (AUC) values calculated for all intensities tested, only bright intensities (>10 lux) or only dim intensities (<0.002 lux) for *O. compressus* (*n* = 5), *N. sammara* (*n* = 4 and 5 for day and night recordings, respectively), and *M. violacea* (*n* = 4). Data are from absolute sensitivity ERGs normalised either to the maximal response ($V_{max}$) or both $V_{max}$ and eye size. Recordings were conducted at day or night. Data are mean ± SEM.

| Stimulus intensity | Normalisation method | Time | *O. compressus* | | *N. sammara* | | *M. violacea* | |
|---|---|---|---|---|---|---|---|---|
| | | | Mean | SEM | Mean | SEM | Mean | SEM |
| *All* | ***$V_{max}$ only*** | *Day* | 5383940 | 2455929 | 4132788 | 489456 | 1632524 | 610014 |
| | | *Night* | 40091 | 21278 | 404635 | 130309 | 1014455 | 187511 |
| | ***Eye size and $V_{max}$*** | *Day* | 547321 | 257105 | 292540 | 36751 | 133899 | 57034 |
| | | *Night* | 4570 | 2577 | 32958 | 11108 | 99261 | 21357 |
| *Bright* | ***$V_{max}$ only*** | *Day* | 5383724 | 2456007 | 4132670 | 489486 | 1631946 | 610147 |
| | | *Night* | 39316 | 21281 | 404108 | 130359 | 1013724 | 187472 |
| | ***Eye size and $V_{max}$*** | *Day* | 547321 | 257106 | 292540 | 36751 | 133899 | 57034 |
| | | *Night* | 4567 | 2577 | 32958 | 11108 | 99261 | 21357 |
| *Dim* | ***$V_{max}$ only*** | *Day* | 0 | 0 | 0 | 0 | 0.0091 | 0.0057 |
| | | *Night* | 0.0052 | 0.00278 | 0.0098 | 0.0053 | 0.013 | 0.0073 |
| | ***Eye size and $V_{max}$*** | *Day* | 0 | 0 | 0 | 0 | 0.00064 | 0.0004 |
| | | *Night* | 0.00058 | 0.0003 | 0.0007 | 0.00038 | 0.00089 | 0.00044 |
